# Selection and Validation of Reference Genes Desirable for Gene Expression Analysis by qRT-PCR on Seed Germination of *Castanea henryi*

**DOI:** 10.1101/2021.01.27.428382

**Authors:** Bin Liu, Yuting Jiang, Ruqiang Lin, Yuanfang Xiong, Shuzhen Jiang, Hui Lian, Xuedie Liu, Zhong-Jian Liu, Shipin Chen

**Author notes:** Corresponding author (Z-J.L.) and (S.C).

## Abstract

Seed germination is the beginning of the plant’s life cycle, and seed biology is one of the most extensively researched areas in plant physiology, however, *Castanea henryi* as an important seed plant, the stable internal reference gene during germination is not clear. In this study, seven candidate genes (TUA, TUB, TIF, UBC, RPL21, RPL30, RPL34) were screened out from transcriptome data, we analyzed the expression of seven candidate reference genes in *C. henryi* at different germination stages with RT–qPCR, and using common algorithms including NormFinder, geNorm and BestKeeper to evaluate the candidate genes stability. The results showed that RPL34 and RPL30 were selected as the most stable genes by NormFinder; TIF was the most stable gene identified by BestKeeper; RPL34 and RPL21 were the most stable genes ranked by geNorm, and TUB was the most unstable gene identified by all of the three software. The RPL34 gene was used as the reference gene, to detected the expression trend of two starch synthetase genes SS1 and SS2 during germination by RT–qPCR, the results of RT–qPCR and transcriptome sequencing were basically consistent, which verified the stability of RPL34 candidate gene. Our result is not only showed functional genes for germination of *C. henryi* seeds and provide useful guidelines for the selection of reliable reference genes for the normalization of RT– qPCR data for germination of seed plants.

## Introduction

Seed germination is the basis of plant formation and is also the strongest period of life activity in all life periods of a plant, not surprisingly that seed biology is one of the most extensively researched areas in plant physiology [1,2]. Real-time fluorescence quantitative PCR (RT–qPCR) is a nucleic acid quantitative technology developed on the basis of qualitative PCR technology. By adding specific fluorescent genes to the PCR reaction system, the expression of the target gene is accurately and quantitatively analyzed [3]. It has the characteristics of quantitative accuracy, high sensitivity, strong specificity and wide applicability, and is widely used in molecular biology, transgenic products, food safety testing, genetics and the research of mining new genes and their functions [4–7]. However, the relative quantitative results of RT– qPCR are affected by the quantity and quality of RNA, primer specificity, PCR reaction conditions, etc. [8–10]. To monitor the process of RT–qPCR, it is very important to select appropriate internal reference genes. Internal reference gene refers to the gene used as internal reference in gene research, usually housekeeping gene, which is involved in the process necessary for cell survival and expressed at a relatively stable and constant level. It needs to be corrected by internal reference gene in detecting the change of target gene expression level, and its stability has a decisive impact on the accuracy of RT–qPCR results [7,11,12]. The stable expression of the internal reference gene should be relatively stable in different tissues and organs, different environments and different stress body conditions [12–14]. In the process of plant RT–qPCR analysis, some relatively stable housekeeping genes, such as tubulin (TUB), actin (ACT), ubiquitin conjugating enzyme (UBC), transcription initiation factors (TIF), and ribosomal protein (RP), were selected as the internal reference genes.

Fagaceae is the main tree species of tropical, subtropical and temperate forests in the northern hemisphere, and also the most important dominant species of angiosperms. Many Fagaceae plants are important economic plants and dual-use plants [15,16]. *Castanea henryi* belongs to the Castanea of Fagaceae. Castanea is widely distributed between East Asia and East North America, and is used to produce wood, tanning agents and food [17,18]. *C. henryi* has important economic value, including starch, soluble sugar, protein, lipids and 18 kinds of amino acids, 8 of which are necessary for human body, and its nutritional value is higher than that of flour, rice and potatoes [19–21], which can not only be used as staple food [22], but also as traditional Chinese medicine [23]. Actin was selected as the internal reference gene [24,25] in the study of cloning related genes of starch branching enzyme and expression related genes of somatic embryogenesis in *Castanea mollissima*. Chen et al. [26] analyzed the expression stability of EF1α, TUA, TUB, UBQ, 18S rRNA and actin in different tissues and organs of *C. mollissima*. Germination is an important biological process in the growth and development, however, the genes stably expressed in different stages of the germination of *C. henryi* seeds have not been discussed. In this study, the expression levels of 7 candidate genes were analyzed by RT–qPCR using 4 seeds in different stages of the germination of *C. henryi*. Three software were used to evaluate the expression stability of candidate internal reference genes, and selected the internal reference genes stably expressed during the germination. Combining with the data of transcriptome sequencing, the expression level of two starch synthesis genes in the process of germination of *C. henryi* was analyzed and compared to further verify the stability of the internal reference genes and provide theoretical basis for the molecular mechanism of germination of *C. henryi*.

## Results

### Candidate gene and RNA quality

The average value, standard deviation and coefficient of variation were calculated according to the expression data of internal reference genes in four stages of germination of *C. henryi*. Through the comparison of variation coefficients, seven internal genes were screened out: TUA (tubulin alpha),TUB (tubulin beta),TIF (translation initiation factor 4A),UBC (ubiquitin-conjugating enzyme E2D), RPL21 (large subunit ribosomal protein L21e), RPL30 (large subunit ribosomal protein L30e) and RPL34 (large subunit ribosomal protein L34e) (Table 1).Primer premier 5.0 was used to design primers, and the results are shown in Table 2. The results of agarose gel electrophoresis showed that the total RNA of the four periods had clear and intact bands of 28S and 18S, and there was no tail phenomenon in the band, indicating that total RNA had no degradation or degradation and had good integrity; the total RNA was not degraded (Figure 1). The results of ultramicro spectrophotometer show that the a260/A280 value is between 1.85–2.20, and the concentration is between 250–800 ng/μL, it shows that the RNA extracted is of high purity, free of protein and inorganic salt pollution, which can meet the needs of subsequent experiments (Table 3).

**Table 1.** Information of seven candidate reference genes

| Gene | Gene ID | Gene annotation | Mean/FPKM | SD | CV |
| --- | --- | --- | --- | --- | --- |
| TUA | Che002914 | tubulin alpha | 15.40 | 3.55 | 0.231 |
| TUB | Che010804 | tubulin beta | 85.80 | 25.27 | 0.294 |
| TIF | Che015192 | translation initiation factor 4A | 218.38 | 12.00 | 0.055 |
| UBC | Che026117 | ubiquitin-conjugating enzyme E2 D | 301.11 | 25.86 | 0.086 |
| RPL21 | Che002638 | large subunit ribosomal protein L21e | 680.30 | 22.62 | 0.033 |
| RPL30 | Che007164 | large subunit ribosomal protein L30e | 469.41 | 13.68 | 0.029 |
| RPL34 | Che034253 | large subunit ribosomal protein L34e | 162.88 | 5.62 | 0.035 |

**Table 2.**
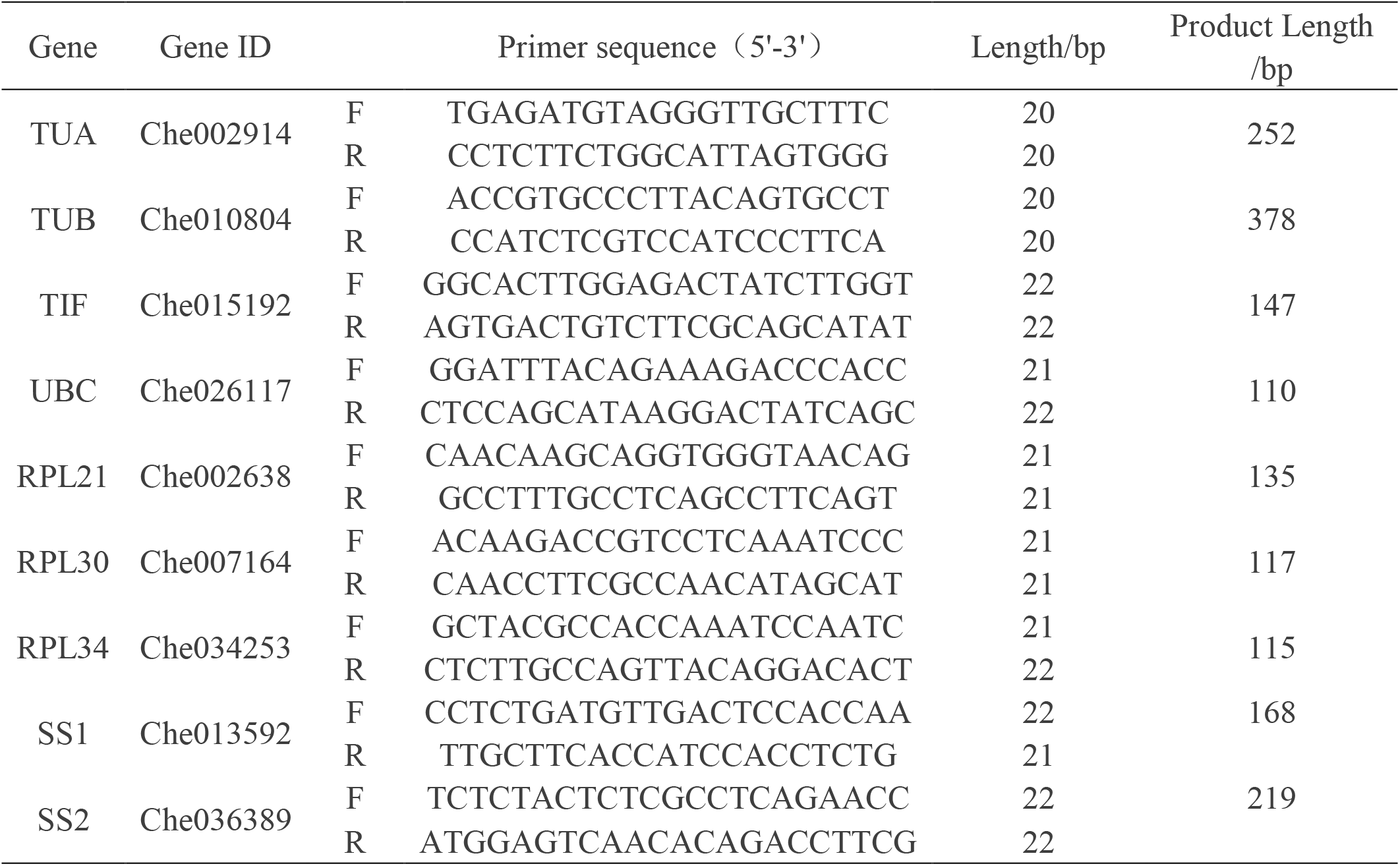
Primer sequences of the candidate reference genes and target genes

**Table 3.**
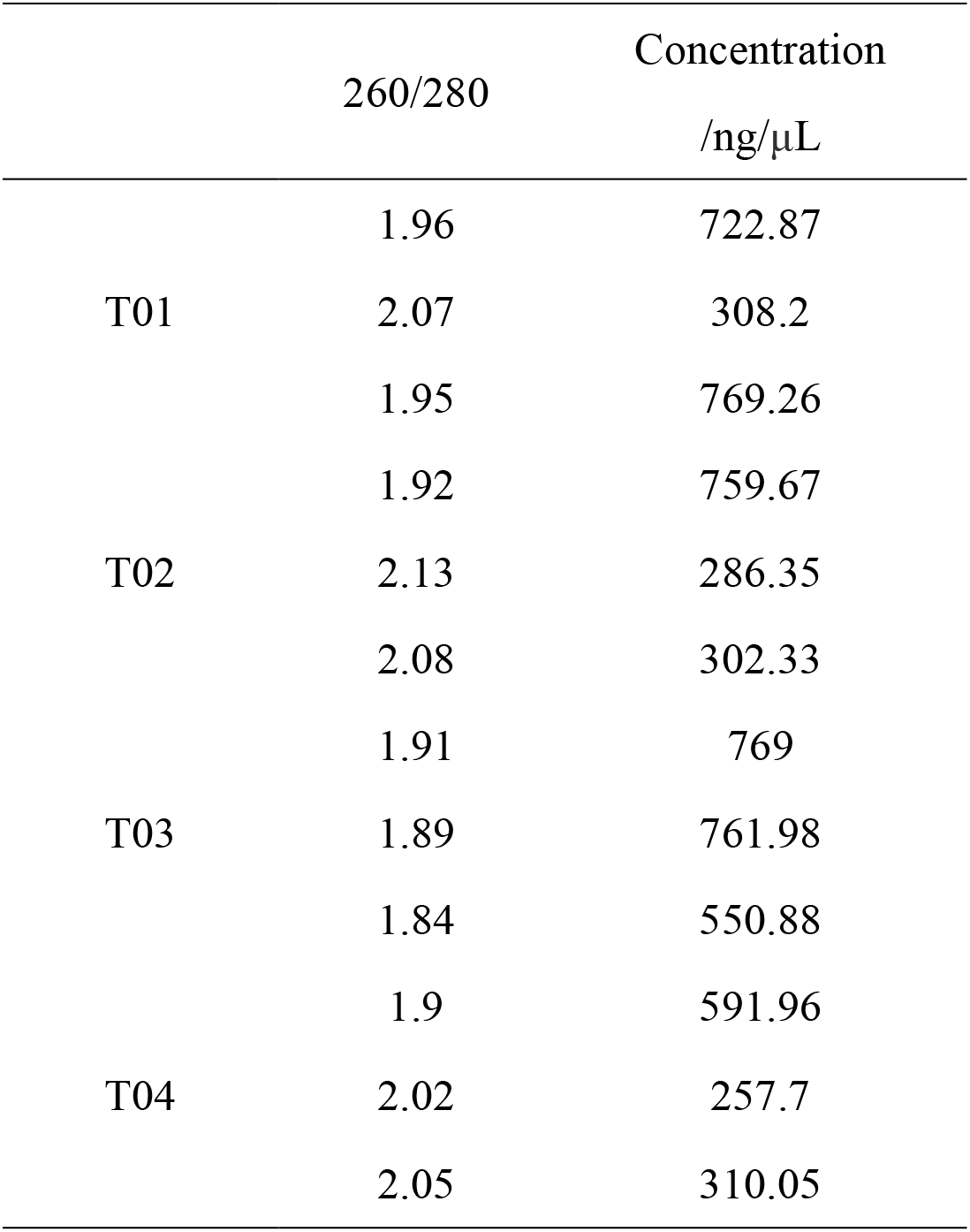
The quality of total RNA in four periods

**Figure 1.**
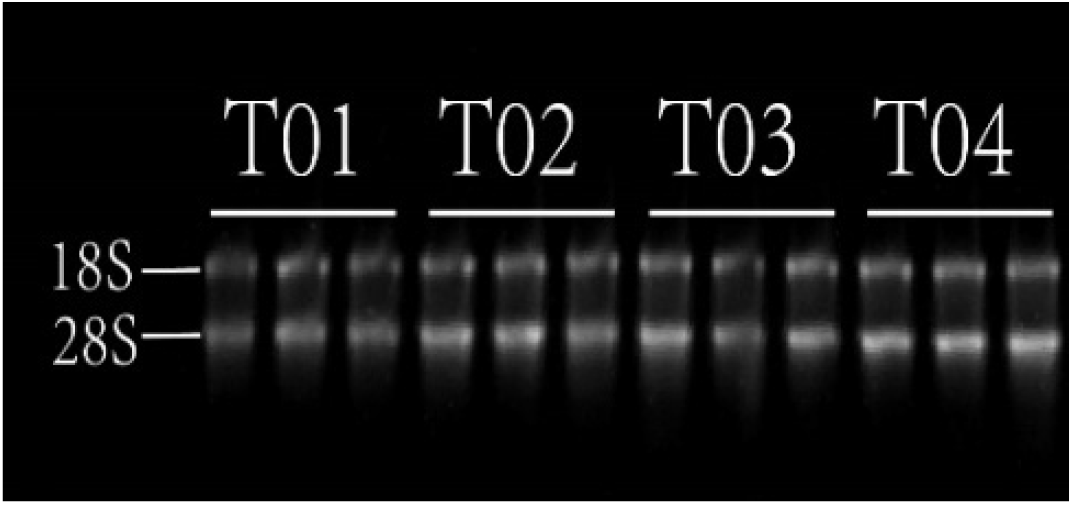
The agarose gel electrophoresis of total RNA.

### Candidate internal reference genes amplification efficiency and linear analysis

The amplification efficiency and linear relationship of the internal reference genes were calculated by RT–qPCR with the mixture of four stages of chestnut kernel cDNA as template, diluted 5 times and diluted five times into four gradients. The results showed that the dissolution curves of the seven internal reference genes were single signal peaks, there was no obvious heteropeak before TM (Figure 2), and the amplification efficiency was between 90% and 105% (Table 4), indicating that the amplification specificity of each pair of primers was good, there was no primer dimer and non-specific amplification in the amplification process, which could be used for subsequent RT–qPCR analysis.

**Table 4.**
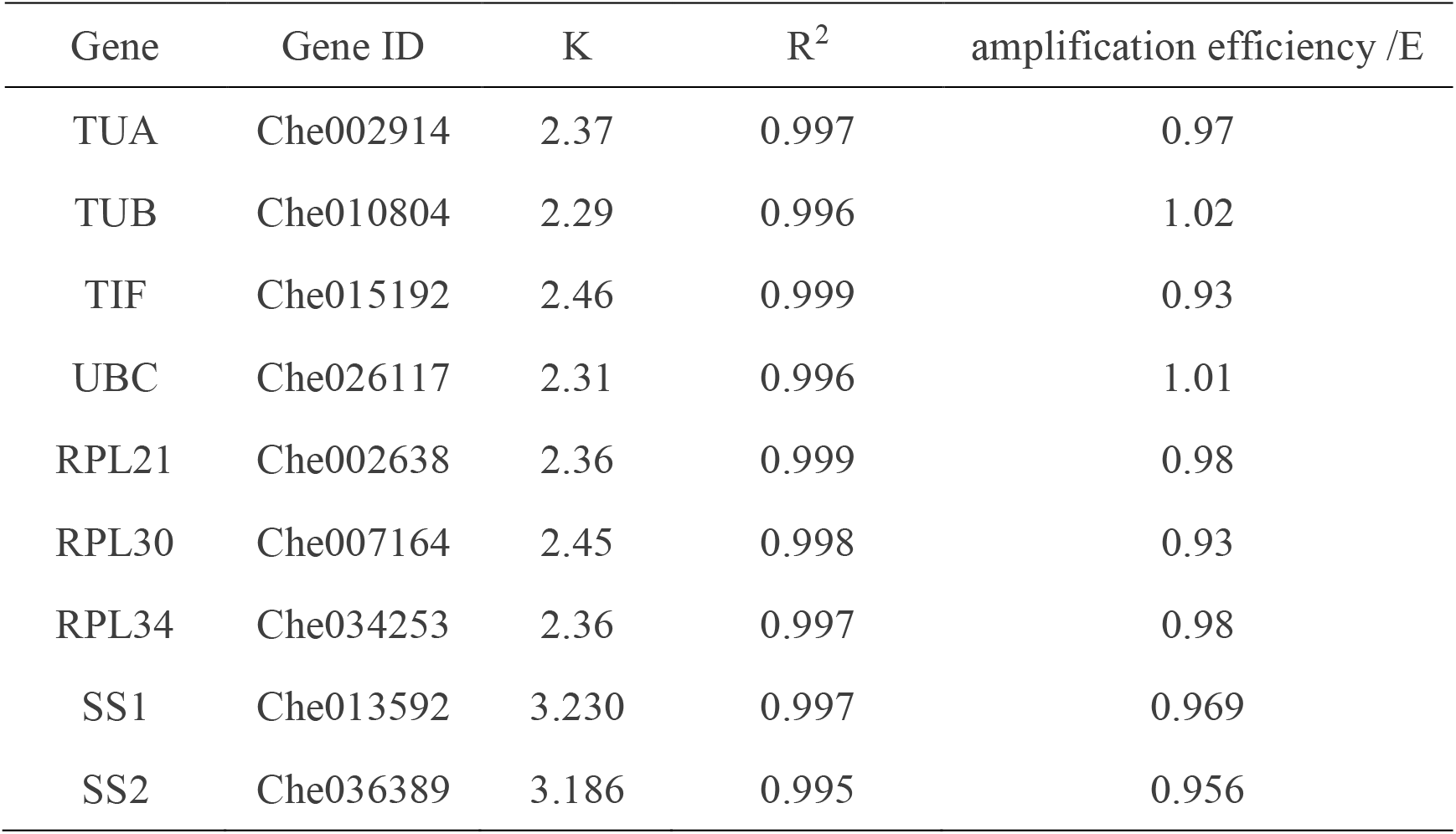
The amplification efficiency of candidate internal reference genes primers and target gene primers

| Gene | Gene ID | K | R<sup>2</sup> | amplification efficiency /E |
| --- | --- | --- | --- | --- |
| TUA | Che002914 | 2.37 | 0.997 | 0.97 |
| TUB | Che010804 | 2.29 | 0.996 | 1.02 |
| TIF | Che015192 | 2.46 | 0.999 | 0.93 |
| UBC | Che026117 | 2.31 | 0.996 | 1.01 |
| RPL21 | Che002638 | 2.36 | 0.999 | 0.98 |
| RPL30 | Che007164 | 2.45 | 0.998 | 0.93 |
| RPL34 | Che034253 | 2.36 | 0.997 | 0.98 |
| SS1 | Che013592 | 3.230 | 0.997 | 0.969 |
| SS2 | Che036389 | 3.186 | 0.995 | 0.956 |

**Figure 2.**
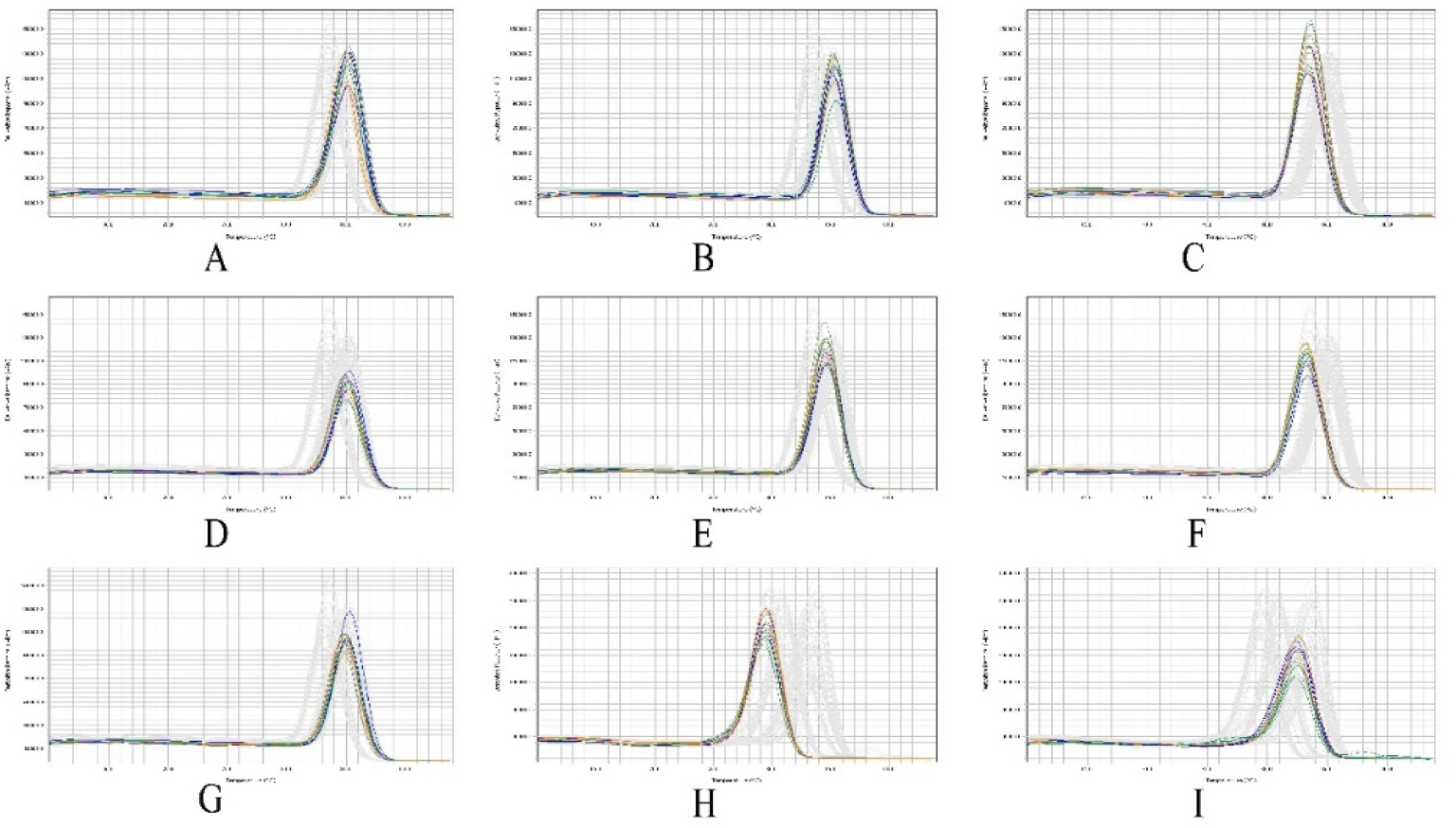
Melting curves of the candidate reference genes and target genes.

### Analysis of stable expression of candidate internal reference genes

The results of CT value analysis showed that the 7 candidate genes had some changes in different stages of germination of *C. henryi*, and the trend was similar. The lower CT value of UBC indicates the higher expression abundance, and the higher CT value of tub indicates the lower expression level of tub. It can be inferred that RPL21, RPL30, RPL34, TIF and UBC can be used as candidate genes. According to the analysis of BestKeeper software, the expression stability of 7 candidate genes is TIF > TUA > RPL21 > RPL34 > RPL30 > UBC > TUB (Figure 3 and Table 5), from which it can be seen that TIF is the most stable one among the seven candidate internal reference genes. According to the analysis results of GeNorm software, the expression stability of the seven candidate internal reference genes is RPL34 / RPL21 > TIF > RPL30 > UBC > TUA > TUB (Figure 3 and Table 5), which shows that RPL34 and RPL21 are better than other candidate internal reference genes in terms of stability, combined with standard deviation According to the analysis results, V_2/3_ is less than 1.5, so the combination of RPL34 and RPL21 can be selected; NormFinder software analysis results show that the expression stability of the seven candidate genes is RPL34 / RPL30 > UBC > TIF > RPL21 > TUA > TUB (Figure 3 and Table 5). The results showed that RPL34 was stable and could be selected as the best internal reference gene for gene expression analysis during seed germination of *C. henryi*.

**Table 5.** The evaluation of candidate reference genes stability from Bestkeeper,GeNorm and Normfinder

|  | Bestkeeper |  | GeNorm |  | Normfinder |  |
| --- | --- | --- | --- | --- | --- | --- |
|  | SD | Rank | M | Rank | S | Rank |
| UBC | 1.50 | 6 | 0.5614 | 4 | 0.209654 | 2 |
| RPL21 | 1.07 | 3 | 0.4159 | 1 | 0.36629 | 4 |
| RPL30 | 1.49 | 5 | 0.5173 | 3 | 0.131927 | 1 |
| RPL34 | 1.22 | 4 | 0.4159 | 1 | 0.131927 | 1 |
| TIF | 0.94 | 1 | 0.435 | 2 | 0.349297 | 3 |
| TUA | 0.99 | 2 | 0.6238 | 5 | 0.587422 | 5 |
| TUB | 2.44 | 7 | 0.9577 | 6 | 1.216812 | 6 |

**Figure 3.**
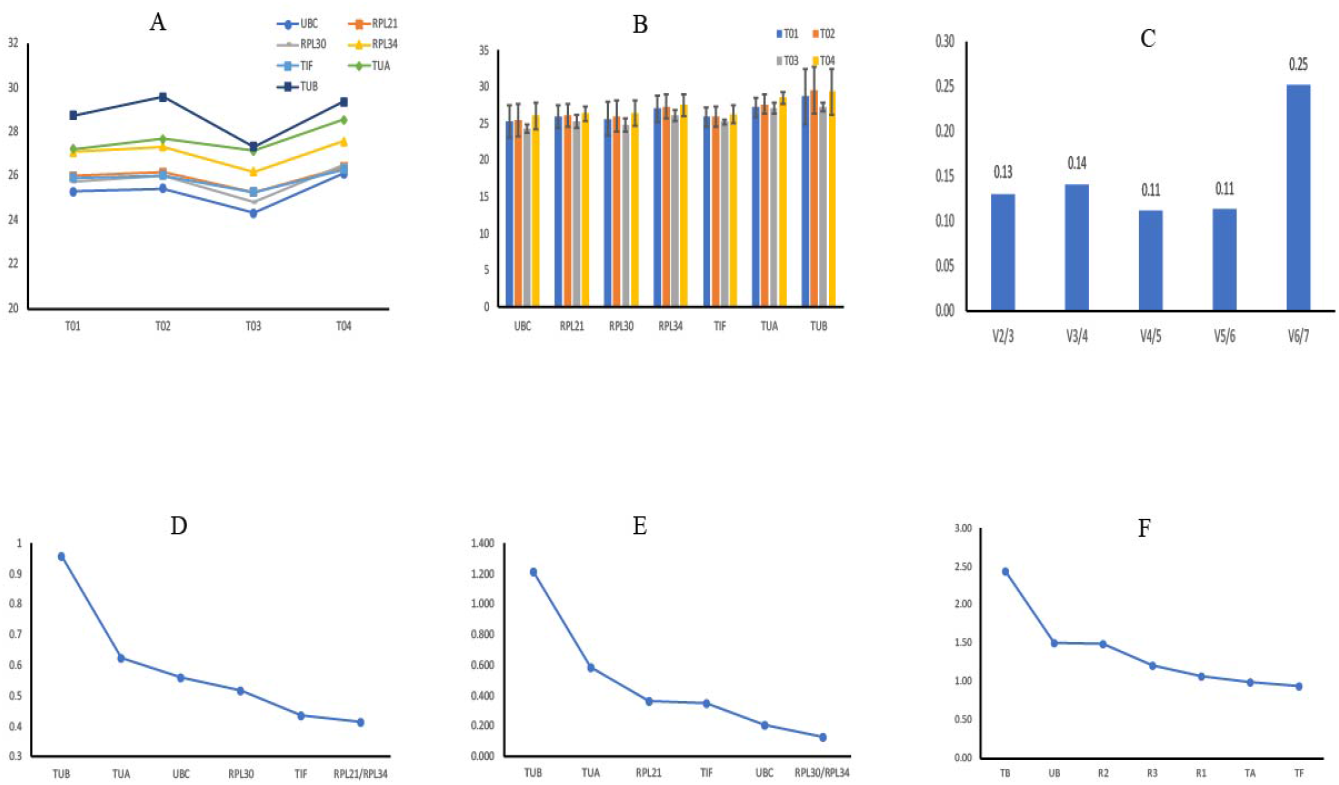
The evaluation of candidate reference genes stability from Ct value, GeNorm,Normfinder and Bestkeeper.

### Validation of RPL34 gene

The expression level of two starch synthetase genes (SS1, SS2) in the germination of *C. henryi* was analyzed by RT–qPCR with RPL34 as the internal reference, and compared with the transcriptome data. The results showed that in the transcriptome data, SS1 and SS2 decreased sharply from T01 to T02, while in T02 to T04, the expression of SS1 continued to increase, while the expression of SS2 changed less. RPL34 was used as the internal reference gene for real-time fluorescence quantitative detection. The expression trend of SS1 and SS2 genes was basically consistent with the results of transcriptome sequencing (Figure 4). The results showed that RPL34 screened from 7 candidate genes was stable in the germination process of *C. henryi*, and it was suitable to be used as an internal reference gene.

**Figure 4.**
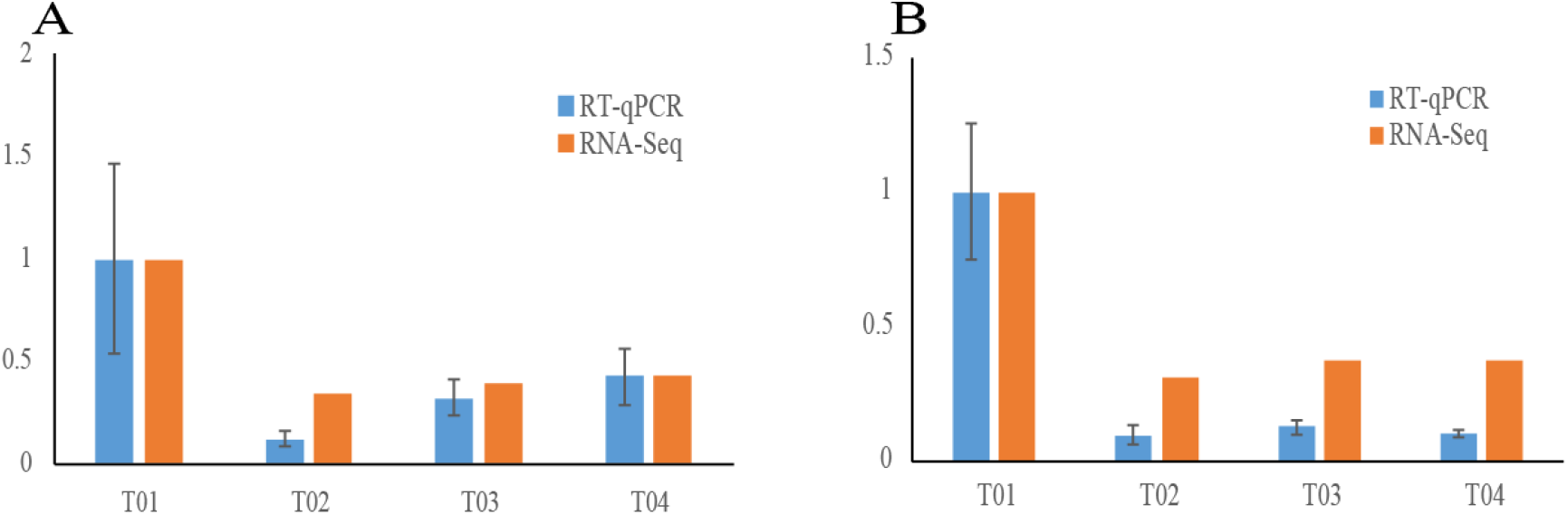
The result of target gene’s RT–qPCR and transcriptome sequencing.

## Discussion

RT–qPCR technology is widely used in the study of gene expression verification. The selection of stable internal reference genes is an important condition to ensure the accuracy of the results [7,27–29]. Ideally, the expression amount of the internal reference gene is relatively stable in different development stages, different tissues and different stress conditions of the sample, but the stability is not absolute. At present, no single fixed internal reference gene can meet all experimental requirements, so it is necessary to select a relatively stable internal reference gene according to the actual experimental conditions [30–32]. It was found that ubiquitin converging enzyme (UBC), actin (ACT) and phospholipase A22 (PLA) were the most stable internal reference genes during *Plukenetia volubilis* seedling and flower development, while UBC *30S ribosomal protein S13*(RPS13)and RNA polymerase II subunit (RPII) were the most stable during seed development [33]. The conclusion of research on *Xanthoceras sorbifolia* is that UBQ and EIF-4α are the most stable in the period of sex differentiation, while EF-1α and EF-1α are the most stable in different organs [34]. Under different stress conditions, the stable expression genes of *Caragana nipponica* were screened, it was found that the combination of UNK2, SAND family protein (SAND) and elongation factor 1-α (EF–1α) was the most stable under salt stress, the combination of TIP41-like family protein (TIP41) and protein photosphatase 2A (PP2A) was the most stable under PEG treatment, and the combination of sand, PP2A and TIP41 was the most stable under heat stress, while the combination of SAND and EF-1 α was the most stable under cold stress [35]. In order to study the expression of related genes in the process of somatic cell development and fruit formation of *C. mollissima*, actin was selected as the internal reference gene for the experiment, GAPDH was selected as the internal reference gene in the study of oxalate oxidase (oxo) gene expression in the process of leaf necrosis caused by *Cryptosporidium parasitica* in *C. dentata*, and Chen found that actin and EF1 α had the best stability in different tissues and organs of *C. mollissima*, [24–26,36].

Seed germination is an important stage in the life cycle of plants, it is one of the urgent problems to screen the relatively stable internal reference genes during germination of *C. henryi*. In this paper, based on the annotation analysis of transcriptome sequencing data and the preliminary comparison of the expression stability of several kinds of housekeeping genes, seven candidate genes (TUA, TUB, TIF, UBC, RPL21, RPL30 and RPL34) were selected. Combined with the real-time fluorescence quantitative technology, three software geNorm, NormFinder and BestKeeper were used for further stability analysis and comparison. In the research process of *Amygdalus persica, Lolium multiflorum* and *Ping’ou Hybrid Hazelnut* (*C*.*heterophylla* × *C*.*avellana*), the internal reference genes were screened based on the transcriptome sequencing data and further combined with the analysis software [37-39]. GeNorm, NormFinder and BestKeeper are the most commonly used analysis software in the current screening of internal reference genes. The evaluation of stability of internal reference genes by the three is based on different algorithms and evaluation indicators, which may lead to inconsistent analysis results. Among them, the analysis results of geNorm and NormFind based on the relative expression Q value are relatively close, while the results of the BestKeeper analysis may be slightly different from the first two which based on CT value [40,41]. In the screening of young bulb internal reference genes in different tissues, hybrids and flower development of Lycoris, it was found that the analysis results of geNorm and NormFinder software were close, while the analysis results of BestKeeper software were significantly different [42]. In addition, similar situations occurred in the internal reference screening of *Siraitia grosvenorii, Ampelopsis grossedentata* and Citrus [43**–** 45].

In this study, the expression stability of 7 candidate genes in the seeds of *C. henryi* during germination was analyzed and compared. The results of geNorm showed that RPL34 and RPL21 had the best stability, the results of NormFinder showed that RPL34 and RPL30 had the best stability, while the BestKeeper’s analysis results show that TIF has the best stability and RPL34 has the general stability. TUB had the worst stability in the three analysis results. Ribosomal protein (RP) is the main component of ribosome, which plays an important role in protein biosynthesis in cells. Ribosomal protein includes large subunit ribosomal protein and small subunit ribosomal protein [45]. Comparing the stability of ten kinds of internal reference genes in different development stages of peanut seed samples and different tissues, it was found that the stability of alcohol dehydrogenase class III (ADH3) was the best, followed by 60S ribosomal protein L7 (60s) [47]. TUB gene plays an important role in maintaining cell structure. It is used as a reliable internal reference gene in switchgrass and peach research. However, in this study, the stability of TUB gene is poor in the germination process of *C. henryi*. This result is similar to that of potato and Soybean [48–51]. Based on the above results, RPL34 has the best stability in the germination process of *C. henryi*, which is suitable for studying the molecular mechanism of starch metabolism during seed germination. The stability of RPL 34 during the germination of *C. henryi* was further confirmed by the expression of SS1 and SS2.

## Conclusion

In this study, through the analysis of the expression stability of seven candidate genes (TUA, TUB, TIF, UBC, RPL 21, RPL30 and RPL34) in the germination process of *C. henryi*, RPL34 gene was screened out which was stably expressed in different stages of the germination process, and the expression stability of TUB was poor, which was not suitable to be the internal reference gene in the germination process of *C. henryi*. RPL34 gene was used as the reference gene to analyze the relative expression of SS1 and SS2 genes in different germination stages of *C. henryi*. The results of RT–qPCR and transcriptome sequencing were basically the same: the expression of SS1 and SS2 was the highest in the first stage, decreased rapidly in the second stage, and changed little in the latter two stages. In order to improve the accuracy of RT–qPCR experiment in the germination process of *C. henryi*, a suitable internal reference gene was screened.

## Materials and methods

### Plant materials

The *C. henryi* variety “Youzhen” was planted in the research center of Youcha, Fujian agricultural and Forestry University. From December 2019 to January 2020, samples were taken every five days. According to the difference between the time interval and kernel morphology, four periods were selected: 0 d, 10 d, 20 d and 30 d for the study, and the transcriptome sequencing was carried out. During sampling, the seed shell and exotesta were peeled off quickly. The whole plant was washed and dried with pure water, and then put into a 50 ml centrifuge tube, frozen with liquid nitrogen, and stored in an ultra-low temperature refrigerator at -80 °C for future use. Three biological repeats were taken from each period.

### RNA extraction and cDNA synthesis

Total RNA from the seeds was isolated using the RNAprep Pure Plant Kit (Polysaccharides and Polyphenolics, Tiangen, Beijng, China) according to the manufacturer’s instructions. The integrity of the total RNA sample was verified by 1% agarose gel electrophoresis. The purity and concentration of the samples were detected by NanoDrop 2000c ultramicro spectrophotometer (Thermo Scientific, USA), and the A260 /A280 ratio and purity were determined. Three duplicates of total RNA extracts were reverse transcribed for first-strand cDNA synthesis for RT–qPCR using the TransScript® All-in-One First-Strand cDNA Synthesis SuperMix for qPCR (One-Step gDNA Removal) (TransGen Biotech, Beijing, China). Finally, it was stored in a refrigerator at -20°C.

### Selection of internal reference gene and primer design

According to the traditional housekeeping genes and the expression amount of these genes in the transcriptome data of four periods during the germination of *C. henryi*, seven genes with stable expression amount were selected as candidate internal reference genes, namely tub, TUA, RPL 21, RPL30, RPL 34, TIF and UBC. The CDS sequences of 7 genes were introduced into primer 5 design primers. The parameters were set as follows: annealing temperature 60 °C, primer length 20-22 bp. Primer blast in NBCI was used to detect the specificity of primers. The primer sequence was compared with transcriptome data to detect the specificity of primers.

### Amplification efficiency of primers for candidate internal reference genes

Take 2 μL from each sample and dilute it five times to get the mixed sample. Take 20 μL mixed sample and dilute it five times to four gradients. That is to say, the final concentration of standard curve template is 24 ng/mL, 4.8 ng/mL, 0.96 ng/mL and 0.08 ng/mL respectively, and each reaction is set with three repeats. The RT–qPCR, performed by TransStart® Tip Green qPCR SuperMix (TransGen Biotech, Beijing, China) with the total system was 20 μL, including 2 μL cDNA. The RT–qPCR conditions followed the manufacturer’s instructions. The primers were designed using Primer 6.0 software. The thermal cycling protocol was 30 s at 94 °C, then 40 cycles of 94 °C for 5 s and 60 °C for 30 s for annealing and extension. Standard curve, slope (k), correlation coefficient (R^2^) and amplification efficiency (E) were obtained by data analysis with CT value.

### RT–qPCR analysis and stability evaluation

According to the RT–qPCR reaction system in 1.4, according to the CT value, we analyzed and compared the expression stability of seven candidate genes in different periods by using the software of geNorm, NormFinder and BestKeeper [52–55]. Cycle threshold (CT) in RT–qPCR was converted according to software requirements. Georm software determines the stability according to the average expression stability value (M). The smaller the m value is, the more stable it is. At the same time, it determines the optimal number of internal reference genes according to V_N_/V_N+1_. NormFinder software sequenced the candidate internal reference genes by the stability of gene expression (s). The genes with the lowest s value had the best stability. The best keeper software mainly compares the standard deviation (SD) and Pearson product motion correlation coefficient (R). If the SD value is less than 1, the closer the R value is to one, the more stable the internal reference gene is. The expression level of the selected target gene in the seed germination process of *C. henryi* was quantitatively analyzed and the validity of the internal reference gene was verified. The relative expression of the target gene was calculated by 2^-ΔΔCt^ method, and each sample was repeated three times.

## Author Contributions

Conceptualization, B.L., S.C., and Z-J.L.; methodology, B.L., S.C., and Z-J.L.; software, B.L.; validation, B.L., Y.J., R.L., and Y.X.; formal analysis, B.L.; investigation, B.L., Y.J., R.L., H.L., X.L. and S.J.; resources, B.L., S.C., and Z-J.L.; data curation, B.L., Y.J., R.L., H.L., X.L. and Y.X.; writing—original draft preparation, B.L., Y.J. R.L.,; writing—review & editing, B.L., S.C., and Z-J.L.; visualization, B.L.; supervision, S.C, and Z-J.L.; project administration, B.L., S.C., and Z-J.L.; funding acquisition, B.L. Z-J.L. and S.C.

## Acknowledgments

The authors would like to thank the Key Laboratory of National Forestry and Grassland Administration for Orchid Conservation and Utilization for supporting this project.

